# m^1^A58 governs two steps of human initiator-methionine tRNA metabolism: maturation and XRN2-mediated decay

**DOI:** 10.64898/2026.05.28.728624

**Authors:** Kiito Otsubo, Mio Nagura, Makoto Terauchi, Hideki Noguchi, Masato T. Kanemaki, Keita Miyoshi, Kuniaki Saito

**Author notes:** Corresponding authors: Keita Miyoshi, Ph.D., Invertebrate Genetics Laboratory, Department of Chromosome Science, National Institute of Genetics, Research Organization of Information and Systems (ROIS), 1111, Yata, Mishima, Shizuoka, 411-8540, Japan., Kuniaki Saito, Ph.D., Invertebrate Genetics Laboratory, Department of Chromosome Science, National Institute of Genetics, Research Organization of Information and Systems (ROIS), 1111 Yata, Mishima, Shizuoka, 411-8540, Japan. These authors contributed equally to this work (Kiito Otsubo, Mio Nagura).

## Abstract

N¹-methyladenosine at position 58 (m¹A_58_), installed by the TRMT6/TRMT61A complex, is a highly conserved structural modification implicated in tRNA stability and human disease. In yeast, loss of m¹A_58_ disrupts initiator methionine tRNA (tRNAᵢᴹᵉᵗ) metabolism through defects in precursor processing and rapid tRNA decay, but whether these mechanisms operate in human cells has remained unclear. Here, using acute degron-mediated depletion of TRMT6 in human HCT116 cells combined with precursor-resolved tRNA sequencing, we show that m¹A_58_ controls two distinct steps of tRNAᵢᴹᵉᵗ metabolism. TRMT6 depletion selectively reduces mature tRNAᵢᴹᵉᵗ while causing accumulation of precursor species retaining both 5′-leader and 3′-trailer sequences, consistent with impaired maturation. In parallel, the nuclear 5′→3′ exonuclease XRN2 selectively degrades the residual mature hypomodified tRNAᵢᴹᵉᵗ pool without affecting precursor accumulation. Loss of mature tRNAᵢᴹᵉᵗ activates the integrated stress response and impairs proliferation, which is restored by XRN2 co-depletion. Together, our findings identify m¹A_58_ as a coordinator of human initiator-tRNA maturation and surveillance, establishing a two-step pathway that maintains tRNAᵢᴹᵉᵗ homeostasis in human cells.

## Introduction

Transfer RNAs (tRNAs) are the most heavily modified class of cellular RNAs, carrying on average more than ten chemically distinct modifications per molecule across their ∼76-nucleotide length ^1^. Far from being passive ornaments, these modifications are now recognized as active regulators of tRNA folding, stability, codon decoding, aminoacylation, and interactions with the translation machinery ^2–5^. They are deposited and, in some cases, removed by a dedicated set of writer, eraser, and reader enzymes whose dysregulation is increasingly implicated in human disease, ranging from intellectual disability and mitochondrial disorders to cancer ^5–7^.

Among structural-core modifications, N¹-methyladenosine at position 58 (m¹A_58_), located on the T-loop, is one of the most evolutionarily conserved tRNA modifications, present from bacteria to humans ^8,9^. In eukaryotes, m¹A_58_ is deposited by a heterotetrameric methyltransferase complex composed of two TRMT6 (Gcd10/Trm6 in yeast) regulatory subunits and two TRMT61A (Gcd14/Trm61) catalytic subunits, in which TRMT6 is essential for substrate recognition and for the structural integrity of the active complex ^8–11^. The N¹-methyl group on A58 introduces a positive charge under physiological pH, neutralizing the negatively charged ribose-phosphate backbone in the T-loop and stabilizing the reverse Hoogsteen base pair between A58 and T54 that locks the elbow region of the L-shaped tertiary fold ^12,13^. Initiator methionine tRNA (tRNAᵢᴹᵉᵗ) carries several sequence features that distinguish it from elongator tRNAs — most notably an A1–U72 base pair at the acceptor stem and an A20 in the D-loop ^12,14^— but whether these features translate into a greater functional dependence on m¹A_58_ in human cells has not been directly tested. Functionally, tRNAᵢᴹᵉᵗ is the substrate of the eIF2·GTP·Met-tRNAᵢᴹᵉᵗ ternary complex that delivers methionine to the small ribosomal subunit during 5′-cap-dependent translation initiation, and its abundance and integrity are therefore rate-limiting for global protein synthesis and for the integrated stress response (ISR) signaling axis that monitors translation initiation ^15–17^.

In yeast, m¹A_58_ is especially important for tRNAᵢᴹᵉᵗ metabolism *in vivo*. Genetic loss of the yeast TRMT6/TRMT61A orthologs *trm6* (*gcd10*) and *trm61* (*gcd14*) reduces mature tRNA levels but also causes the accumulation of precursor tRNAs — most strikingly precursors of tRNAᵢᴹᵉᵗ that retain their 5′ leader and 3′ trailer sequences — indicating that m¹A_58_ supports an early step of tRNA processing ^8,18,19^. Aberrant pre-tRNAs in yeast are polyadenylated by the nuclear TRAMP complex (Trf4–Air1/2–Mtr4) and degraded by the nuclear exosome ^19,20^. In parallel, hypomodified mature tRNAs in yeast are selectively eliminated by a dedicated pathway termed rapid tRNA decay (RTD), in which the 5′→3′ exonucleases Rat1 and Xrn1 act redundantly to degrade aberrant mature tRNAs ^21–24^. Genetic disruption of Rat1 — the yeast ortholog of mammalian XRN2 — suppresses the loss of tRNAᵢᴹᵉᵗ caused by m¹A_58_ deficiency in trm6/trm61 mutants. In addition, recent work showing that Rat1 and its orthologs act on tRNA-derived RNAs across phylogenetically distant yeast species further supports a conserved role for nuclear 5′→3′ exonucleases in tRNA-related RNA metabolism ^25^. Together, these findings suggest that m¹A_58_ contributes both to precursor maturation and to protection of the mature tRNA pool.

In mammalian cells, depletion of the m¹A writer complex TRMT6/TRMT61A has been associated with proliferation defects in several human cancer cell models, including liver, bladder, colorectal, and thyroid cancers ^26–29^. In rat C6 glioma cells, TRMT6 knockdown reduced RT-qPCR signals for initiator tRNAᵢᴹᵉᵗ, while phenotypic rescue by tRNAᵢᴹᵉᵗ overexpression suggested a functional connection between TRMT6 activity and initiator tRNA–dependent processes ^30^. A separate study has reported that mature tRNAᵢᴹᵉᵗ accumulates in the nucleus of heat-stressed HeLa cells and is targeted there by the 5′→3′ exonucleases XRN1 and XRN2 ^31^; whether an analogous surveillance pathway operates under physiological growth conditions, and whether it is coupled to the loss of TRMT6/TRMT61A-mediated m¹A_58_ modification, has remained unknown. However, it has not been established whether loss of m¹A_58_ modification in human tRNAᵢᴹᵉᵗ primarily affects precursor maturation, promotes selective decay of mature tRNAᵢᴹᵉᵗ, or both. Addressing this question has been challenging because long-term depletion of TRMT6/TRMT61A induces extensive secondary adaptation.

Here, using acute degron-mediated depletion ^32^ of TRMT6 together with RNA sequencing, we show that m¹A_58_ controls two mechanistically separable steps of human tRNAᵢᴹᵉᵗ metabolism. TRMT6 depletion causes accumulation of precursor tRNAᵢᴹᵉᵗ species and reduction of mature tRNAᵢᴹᵉᵗ, while the human exonuclease XRN2 selectively degrades the residual mature hypomodified pool. Acute TRMT6 loss activates the ATF4-mediated integrated stress response and impairs cell proliferation. Co-depletion of XRN2 selectively restores mature tRNAᵢᴹᵉᵗ levels without altering the accumulated precursor pool, and rescues the proliferation defect. These analyses demonstrate that m¹A_58_ governs two molecularly separable steps of tRNAᵢᴹᵉᵗ metabolism: it licenses efficient precursor maturation, and it protects the mature tRNA pool from XRN2-mediated decay in human cells.

## Results

### Acute, degron-mediated TRMT6 depletion abolishes m¹A on cytosolic tRNAs and selectively reduces mature tRNAᵢᴹᵉᵗ

To dissect the immediate consequences of TRMT6 loss on physiological timescales, we established an acute ligand-inducible TRMT6 depletion system using the BromoTag degron platform ^32,33^, enabling near-complete protein depletion within hours while minimizing secondary compensatory effects. Using CRISPR-Cas9-mediated knock-in, we tagged the N-terminus of the endogenous TRMT6 locus in HCT116 cells with a BromoTag preceded by a P2A-blasticidin selection cassette (Fig. 1a). Following blasticidin selection, we screened for clones carrying biallelic homologous recombination at the TRMT6 locus by genomic PCR and isolated two independent BromoTag-TRMT6 knock-in clones (KI#1 and KI#2) and used both in parallel throughout this study to confirm that the observed phenotypes are not clone-specific artifacts of CRISPR editing. To test whether AGB1 treatment induces efficient degradation of BromoTag-TRMT6, we treated BromoTag-TRMT6 cells with AGB1 and observed a near-complete loss of BromoTag-TRMT6 protein within 30 min of induction (Supplementary Fig. 1a), whereas TRMT6 protein levels in parental control cells were unaffected (Fig. 1b). Notably, BromoTag knock-in resulted in an upward shift of the TRMT6 immunoreactive band, with loss of the native TRMT6 band, further supporting successful biallelic targeting of the endogenous TRMT6 locus. We next asked whether acute TRMT6 loss results in a corresponding loss of m¹A on tRNAs. Because the TRMT6/TRMT61A complex is responsible for the predominant source of cytosolic tRNA m¹A_58_ modification in human cells, loss of TRMT6 protein should be reflected in a measurable decrease in bulk m¹A signal within the tRNA fraction ^9,34,35^. RNA blot analysis using an anti-m¹A antibody revealed a progressive decline in m¹A signal within the bulk tRNA size range across the AGB1 time course (0, 12, 24, 48, 72 h) (Fig. 1c, d). These results confirm that acute TRMT6 depletion leads to a rapid and progressive reduction in bulk tRNA m¹A signal, consistent with the established role of the TRMT6/TRMT61A complex as the major cytosolic tRNA m¹A methyltransferase.

**Fig. 1.**
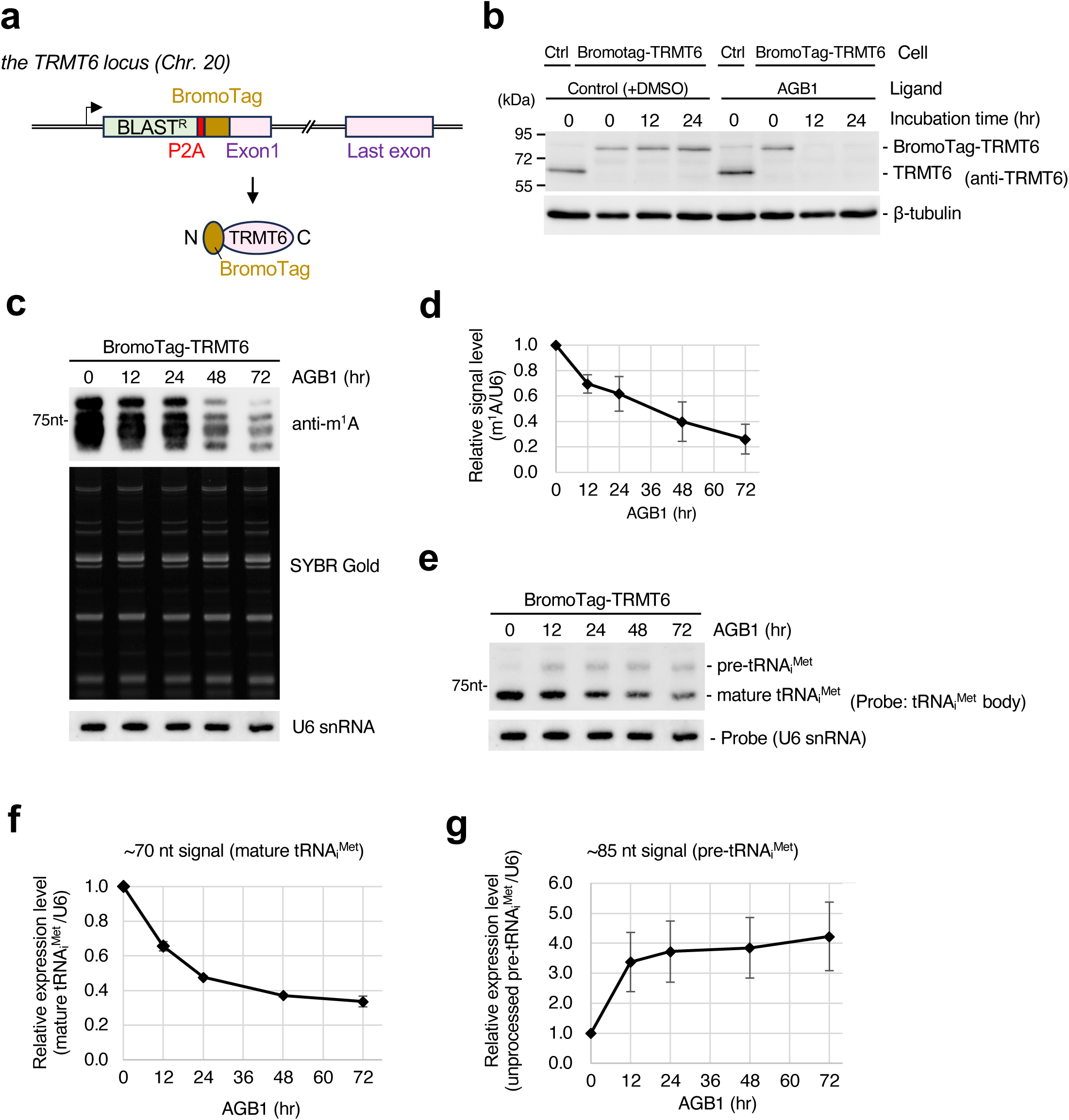
Acute degron-mediated depletion of TRMT6 causes accumulation of precursor tRNAᵢᴹᵉᵗ species and reduction of mature tRNAᵢᴹᵉᵗ. **a,** Schematic of CRISPR/Cas9-mediated knock-in of the BromoTag at the C-terminus of the endogenous TRMT6 locus on chromosome 20. A P2A-blasticidin selection cassette (BLASTR) followed by the BromoTag-encoding sequence was inserted in-frame upstream of the TRMT6 stop codon. **b,** Western blot showing AGB1 (0.5 µM)-induced degradation of BromoTag-TRMT6 in HCT116 BromoTag-TRMT6 cells (KI#1) versus parental control cells, over a 0–24 h time course. β-tubulin, loading control. **c,** Anti-m¹A blot analysis of total RNA from BromoTag-TRMT6 cells treated with AGB1 for the indicated times (0, 12, 24, 48, 72 h). U6 snRNA, loading control. Prior to blotting, total RNA was visualized by SYBR Gold staining to assess RNA integrity and equal loading. **d,** Quantification of bulk tRNA m¹A signal in c, normalized to U6 snRNA. Data are mean ± SD from three biological replicates. **e,** Northern blot of total RNA from BromoTag-TRMT6 cells treated with AGB1 over the same time course as in c, probed with a DIG-labeled oligonucleotide complementary to the conserved body region of tRNAᵢᴹᵉᵗ (Supplementary Fig. 1b). Two species are resolved: mature tRNAᵢᴹᵉᵗ (∼70 nt) and pre-tRNAᵢᴹᵉᵗ (∼85 nt). U6 snRNA, loading control. **f, g,** Quantification of mature tRNAᵢᴹᵉᵗ (f) and pre-tRNAᵢᴹᵉᵗ (g) signal in e, normalized to U6 snRNA. Data are mean ± SD from three biological replicates.

### TRMT6 depletion reduces mature tRNAᵢᴹᵉᵗ and promotes accumulation of pre-tRNAᵢᴹᵉᵗ

To assess the consequences for tRNAᵢᴹᵉᵗ, we performed Northern blotting over the same AGB1 time course using a probe complementary to the conserved tRNA body region shared among all tRNAᵢᴹᵉᵗ isodecoders (Supplementary Fig. 1b). The probe detected two distinct species: a ∼70-nt band corresponding to mature tRNAᵢᴹᵉᵗ (after 5′-leader and 3′-trailer cleavage and 3′-CCA addition) and a slower-migrating ∼85-nt band corresponding to precursor (pre)-tRNAᵢᴹᵉᵗ (Fig. 1e). Over the 48-h depletion time course, mature tRNAᵢᴹᵉᵗ levels progressively declined to ∼40% of control levels, whereas pre-tRNAᵢᴹᵉᵗ accumulated approximately 4-fold (Fig. 1f, g; Supplementary Fig. 1c; d). To our knowledge, endogenous human pre-tRNAᵢᴹᵉᵗ has not previously been clearly resolved by Northern blotting.

### RNA-seq reveals tRNAᵢᴹᵉᵗ-specific accumulation of precursors retaining 5′-leader and 3′-trailer sequences

To define the transcriptome-wide consequences of TRMT6 depletion, we performed RNA-seq on <200-nt RNA fractions isolated from both KI#1 and KI#2 cells treated with AGB1 or DMSO (control). Principal-component analysis (PCA) showed clean separation of AGB1 from DMSO samples on PC1 across both clones (Supplementary Fig. 2a), confirming that TRMT6 depletion is the dominant source of variance in the dataset and that biological replicates cluster tightly. Differential expression analysis of the mature read population identified tRNAᵢᴹᵉᵗ-CAT as the major tRNA species significantly reduced in the mature pool upon TRMT6 depletion in either clone (adjusted *P* < 0.05, log₂ fold-change ≤ −1; Fig. 2a; Supplementary Fig. 2b). Consistent with this result, northern blot analysis detected no substantial changes in the levels of representative non-iMet tRNAs, including tRNA^Lys^-TTT and tRNA^Tyr^-GTA (Supplementary Fig. 2c). Intersection of the significantly downregulated mature tRNAs from KI#1 and KI#2 yielded a single species, tRNAᵢᴹᵉᵗ-CAT, supporting the tRNAᵢᴹᵉᵗ-specificity of the mature-pool loss across independent clones (Fig. 2c; Supplementary Fig. 2d).

**Fig. 2.**
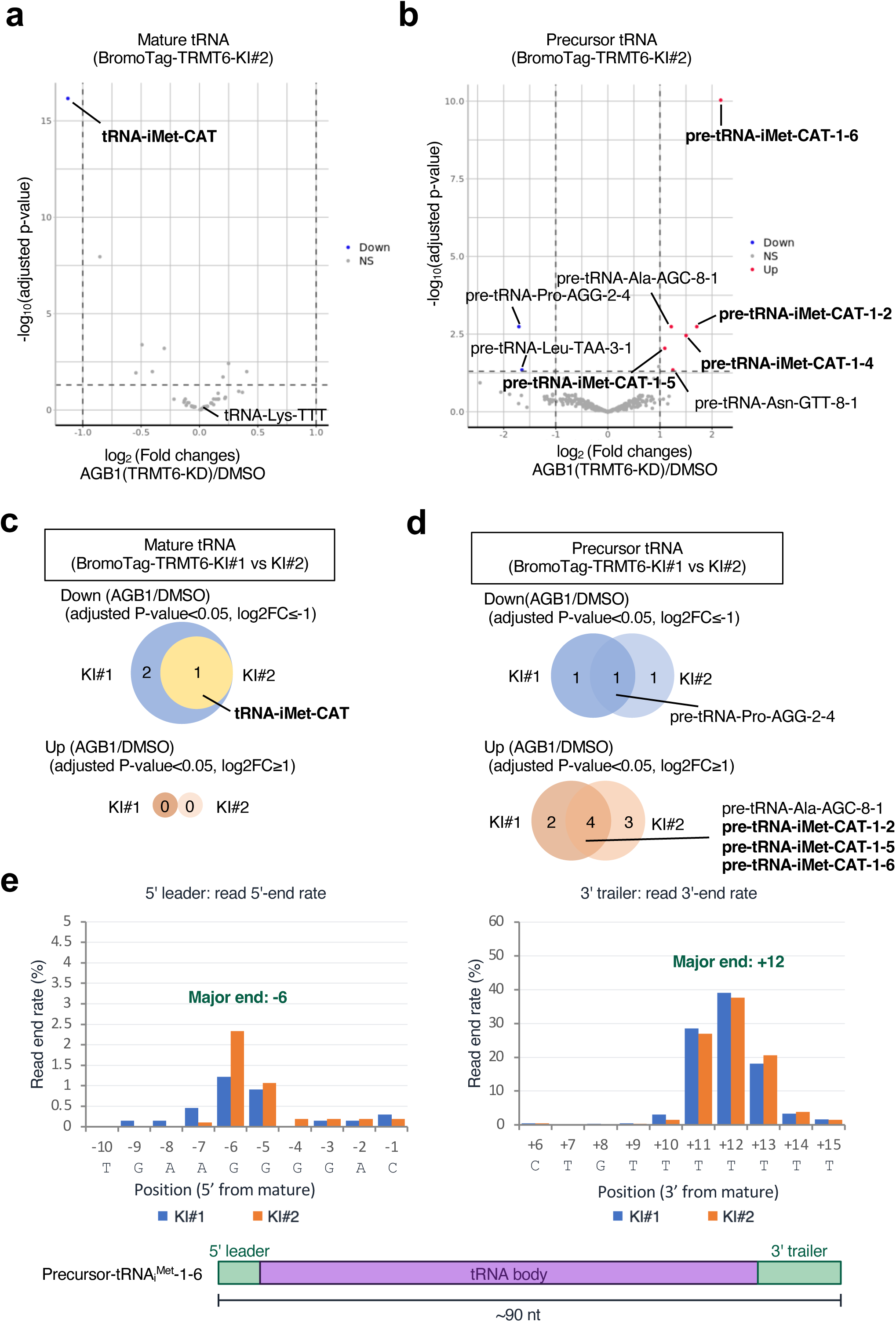
RNA-seq reveals tRNAᵢᴹᵉᵗ-specific accumulation of precursors retaining 5′-leader and 3′-trailer sequences upon TRMT6 depletion. **a**, Volcano plot of tRNA-seq differential expression at the mature-pool, anticodon level in BromoTag-TRMT6 KI#2 cells treated with AGB1 versus DMSO. tRNAs with adjusted *P*-value < 0.05 and |log₂ fold change| ≥ 1 were considered differentially expressed. **b**, Volcano plot of tRNA-seq differential expression for the precursor read population in KI#2 cells. Significance and labeling thresholds as in a; labels containing “tRNA” indicate pre-tRNA at the isodecoder level. **c**, Intersection of significantly down-regulated mature tRNAs (adjusted P < 0.05, log₂FC ≤ −1) between KI#1 and KI#2. The single shared down-regulated species is tRNAᵢᴹᵉᵗ-CAT. **d**, Intersection of significantly up-regulated precursor tRNAs (adjusted *P* < 0.05, log₂FC ≥ 1) between KI#1 and KI#2. Three pre-tRNAᵢᴹᵉᵗ-CAT isodecoders (1-2, 1-5, 1-6) and two non-iMet species (pre-tRNA-Ala-AGC-8-1; pre-tRNA-Pro-AGG-2-4) appear in both clones. **e**, Read 5′-end position (left) and 3′-end position (right) rate around the body of pre-tRNAᵢᴹᵉᵗ-CAT-1-6 in TRMT6-depleted cells. Read end rates were calculated from three TRMT6-knockdown biological replicates as the proportion of reads whose 5′-end fell within the upstream (negative) region or whose 3′-end fell within the downstream (positive) region of the mature tRNA boundaries, relative to the total number of reads mapped to the locus. Green boxes mark the positions with the highest frequencies (5′-end at position −6; 3′-end at position +12). The schematic at the bottom illustrates the inferred pre-tRNAᵢᴹᵉᵗ-CAT-1-6 structure with the 5′-leader and 3′-trailer sequences retained.

We then analyzed the precursor read population. Corresponding to the decrease in mature tRNAᵢᴹᵉᵗ, multiple tRNAᵢᴹᵉᵗ isodecoders (tRNAᵢᴹᵉᵗ-1-2, -1-5, and -1-6) were significantly enriched among accumulated precursors in both clones (adjusted *P* < 0.05, log₂ fold-change ≥1; Fig. 2b, d; Supplementary Fig. 2e). Two additional species, tRNA-Ala-AGC-8-1 and tRNA-Pro-AGG-2-4, also showed modest precursor accumulation and reduction in the two-clone intersection, respectively (Fig. 2d); however, their corresponding mature isodecoders were not significantly affected (Fig. 2a; Supplementary Fig. 2b), and the magnitude of their precursor accumulation was several-fold below that observed for tRNAᵢᴹᵉᵗ isodecoders. We interpret these as minor, possibly indirect effects on the maturation of a small subset of non-tRNAᵢᴹᵉᵗ species, the biological significance of which we did not pursue here. The dominant precursor accumulation arises overwhelmingly from tRNAᵢᴹᵉᵗ isodecoders.

To characterize the maturation status of the accumulated tRNAᵢᴹᵉᵗ precursors and determine which processing step(s) are affected by TRMT6 loss, we mapped read 5′-end and 3′-end positions around the body of tRNAᵢᴹᵉᵗ-CAT-1-6, the isodecoder showing the largest fold change upon TRMT6 loss (Fig. 2b). The major 5′-end mapped 6 nt upstream of the mature 5′-end, and the major 3′-end mapped 12 nt downstream of the mature 3′-end (Fig. 2e), indicating that the accumulated precursors retain both the 5′-leader (normally cleaved by RNase P) and the 3′-trailer (normally cleaved by RNase Z / ELAC2) sequences. End-mapping for the two other accumulated tRNAᵢᴹᵉᵗ isodecoders (tRNAᵢᴹᵉᵗ-CAT-1-2 and -1-5) similarly revealed retention of both 5′-leader and 3′-trailer sequences, with the major end positions varying between isodecoders, reflecting differences in their genomic context (Supplementary Fig. 3a, b). Notably, reads frequently extended into the oligo (T) tract downstream of the tRNA genes, consistent with RNA polymerase III termination signals, indicating that the accumulated precursors appear to retain their original 3′ termini generated by RNA polymerase III transcription termination (Fig. 2e; Supplementary Fig. 3a, b). No clear evidence of TRAMP complex-mediated oligo(A) tail addition was detected. In addition, the pre-tRNA species in samples with and without AGB1 treatment were similar in size, and no apparent accumulation of intermediates carrying only 5′- or 3′-end trimming was observed (data not shown). These observations suggest that the precursor tRNAs themselves accumulated, rather than processing intermediates generated by RNase P- or RNase Z-mediated cleavage at only one end. This interpretation is also consistent with the Northern blot results (Fig. 1e; Supplementary Fig. 1b). Together, these data demonstrate that acute TRMT6 depletion selectively disrupts human tRNAᵢᴹᵉᵗ homeostasis by coupling precursor accumulation with depletion of the mature tRNA pool.

### TRMT6 depletion impairs cell proliferation and activates the integrated stress response

Mature tRNAᵢᴹᵉᵗ, in its aminoacylated form, is delivered to the 40S ribosomal subunit by the eIF2·GTP·Met-tRNAᵢᴹᵉᵗ ternary complex during every protein synthesis initiation event, and is therefore rate-limiting for global cap-dependent translation^17,36^. Cells monitor the availability of this ternary complex through the GCN2-eIF2α-ATF4 branch of the integrated stress response (ISR), which is activated when eIF2 fails to deliver Met-tRNAᵢᴹᵉᵗ at sufficient rate^16^. The Fig. 1 and Fig. 2 phenotypes therefore predict two coupled cellular consequences of acute TRMT6 depletion: a proliferation defect (because translation initiation is impaired) and ATF4 induction (because the ISR is engaged).

AGB1 treatment of BromoTag-TRMT6 cells (KI#1 and KI#2) markedly impaired proliferation over a 3-day time course, whereas parental control cells treated with AGB1 showed no proliferation defect (Fig. 3a). The defect was apparent by day 2 and continued to widen through day 3, consistent with the progressive loss of mature tRNAᵢᴹᵉᵗ over the same time course (Fig. 1f). Both KI#1 and KI#2 showed comparable proliferation defects, again excluding clone-specific artifacts.

**Fig. 3.**
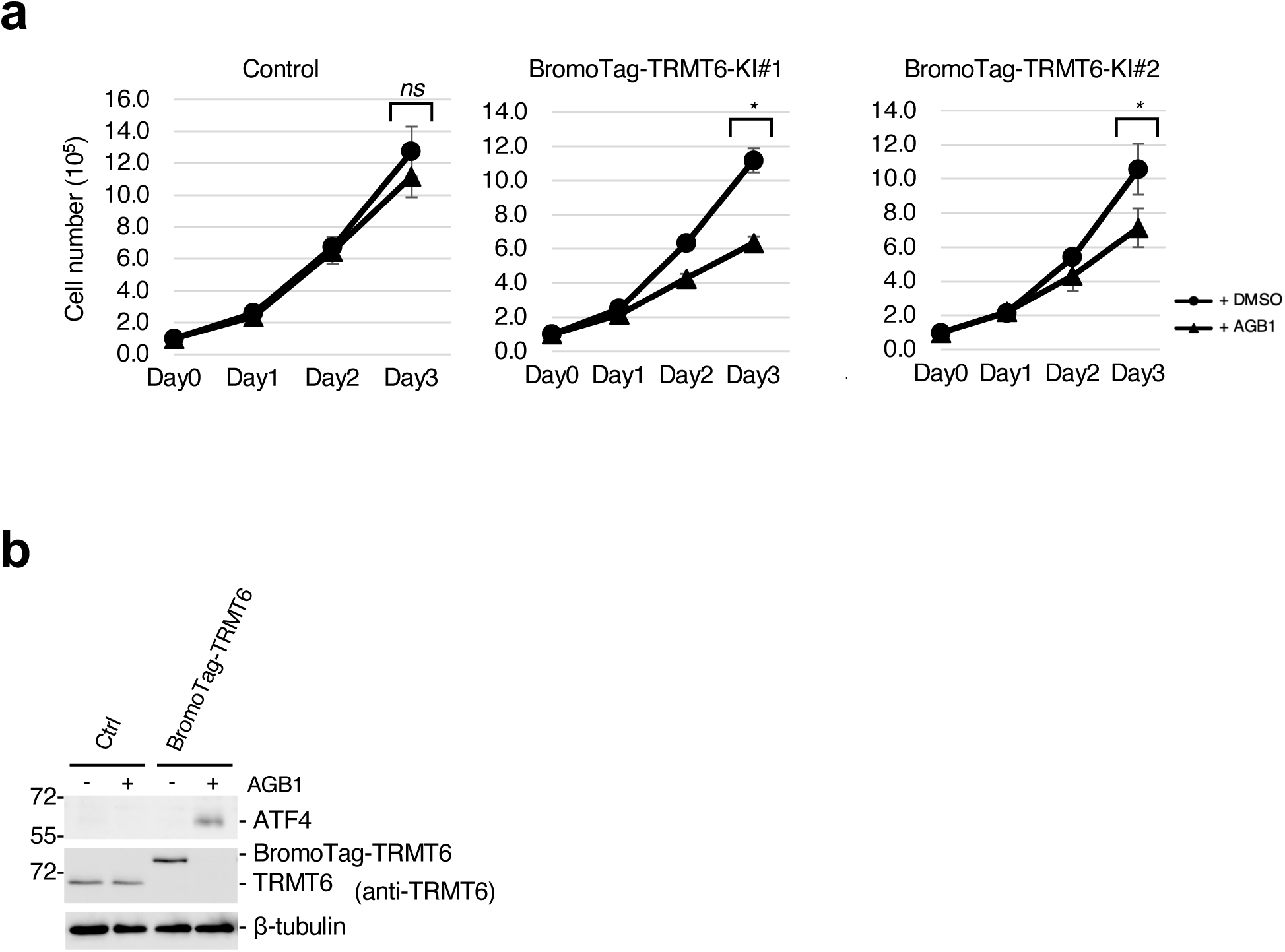
Acute TRMT6 depletion impairs cell proliferation and activates the ATF4-mediated integrated stress response. **a**, Cell proliferation of HCT116 parental control cells (Control) and BromoTag-TRMT6 KI#1 and KI#2 cells treated with DMSO or AGB1 (0.5 µM) over a 3-day time course. Cell number was counted at days 0–3. Data are mean ± SD from three biological replicates. Statistical significance was assessed using Student’s t-test. \**p* < 0.05 for AGB1 vs. DMSO. **b**, Western blot of ATF4 in BromoTag-TRMT6 KI#1 cells and parental control cells treated with AGB1 (or DMSO control). β-tubulin, loading control.

To ask whether the loss of mature tRNAᵢᴹᵉᵗ triggers a translational stress signal, we examined ATF4, the canonical translation-level effector of the ISR. Western blotting revealed robust induction of ATF4 protein in AGB1-treated BromoTag-TRMT6 cells, but not in AGB1-treated parental controls (Fig. 3b). We did not see ATF4 induction in AGB1-treated parental cells (Ctrl), confirming that ATF4 induction depends on TRMT6 loss and not on AGB1 treatment *per se*.

### XRN2 selectively degrades the mature, m¹A_58_-deficient tRNAᵢᴹᵉᵗ

The reduction in mature tRNAᵢᴹᵉᵗ upon TRMT6 loss observed in Fig. 1 could in principle reflect any combination of three mechanisms: (i) impaired maturation alone (the accumulated precursors never enter the mature pool); (ii) accelerated decay of the mature, m¹A_58_-deficient pool by a surveillance nuclease; or (iii) both processes acting in parallel. In yeast, hypomodified tRNAs become substrates of the rapid tRNA decay (RTD) pathway, in which the 5′→3′ exonucleases Xrn1 and Rat1 degrade hypomodified mature tRNAs^21–25^. In mammalian cells, the closest sequence and functional homolog of yeast Rat1 is the nuclear 5′→3′ exonuclease XRN2, and we recently showed in HCT116 cells that XRN2 — but not XRN1 or the nuclear exosome subunits EXOSC10 (Rrp6) — constitutively degrades m⁷G-hypomodified tRNAs (tRNA^Val^-AAC and tRNA^Pro^ isodecoders) under physiological growth conditions, providing the first identification of an RTD-like, XRN2-dependent decay activity in mammals ^37^. Thus, we asked whether XRN2 contributes to the loss of mature tRNAᵢᴹᵉᵗ in TRMT6-depleted human cells. We first took knockdown approach against XRN2 with siRNA in BromoTag-TRMT6 cells and found siRNA-mediated knockdown of XRN2 partially restored the steady-state levels of mature tRNAᵢᴹᵉᵗ levels in AGB1-treated BromoTag-TRMT6 cells (Fig. 4a, b).

**Fig. 4.**
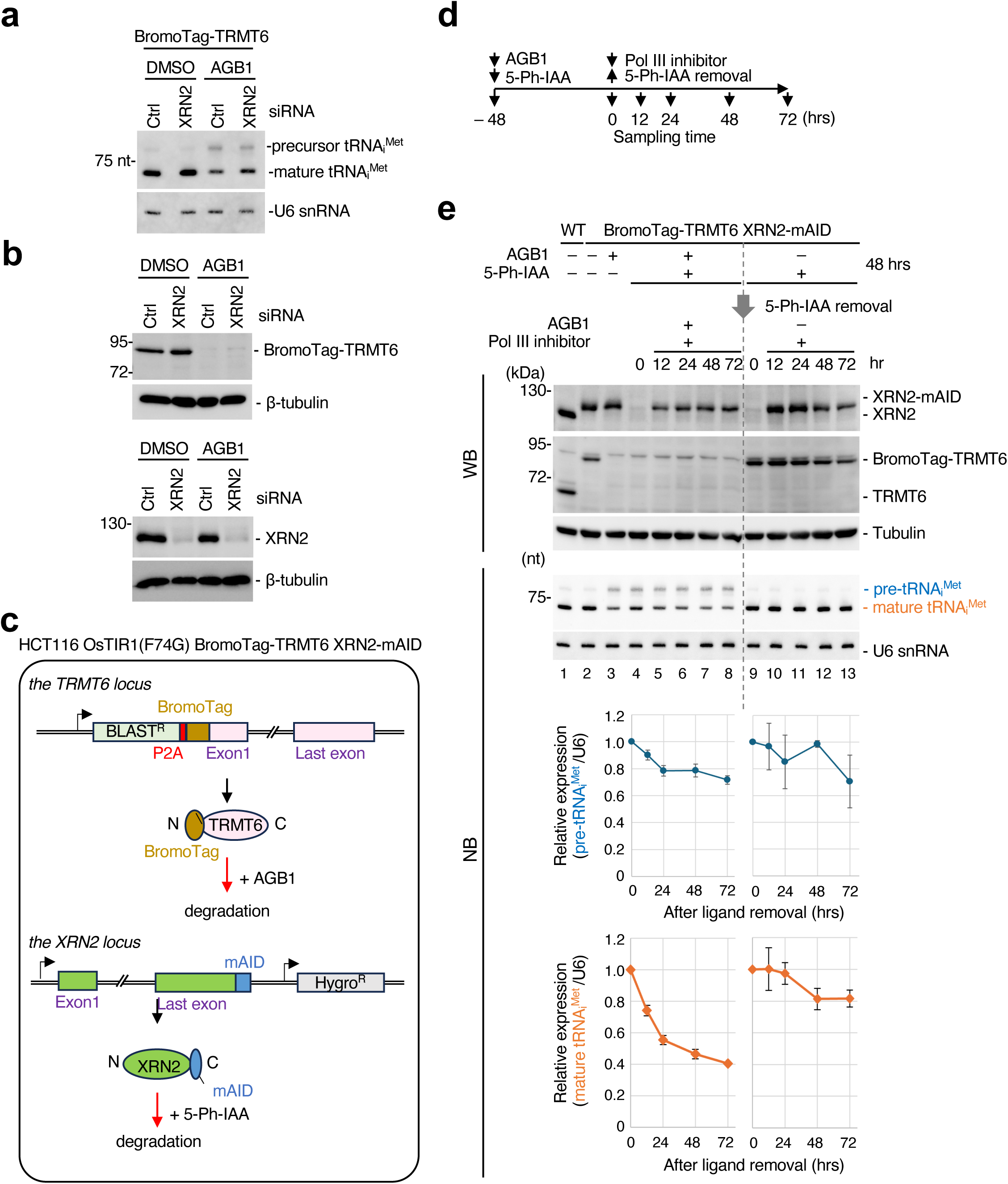
XRN2 selectively degrades the mature, m¹A58-deficient tRNAᵢᴹᵉᵗ. **a**, Northern blot of total RNA from BromoTag-TRMT6 KI#1 cells treated with DMSO or AGB1 (0.5 µM) in combination with non-targeting control siRNA (siCtrl) or siRNA targeting XRN2 (siXRN2) at 10 nM for 48 h. The probe detects mature tRNAᵢᴹᵉᵗ and pre-tRNAᵢᴹᵉᵗ; U6 snRNA, loading control. **b**, Western blot showing BromoTag-TRMT6, XRN2, and β-tubulin levels. **c**, Schematic of the double-degron cell line. BromoTag was knocked in at the N-terminus of TRMT6 (BSD-P2A-BromoTag cassette), and mAID was knocked in at the C-terminus of XRN2 (mAID–HygroR cassette) in the HCT116 OsTIR1(F74G) parental background. AGB1 induces BromoTag-TRMT6 degradation; 5-Ph-IAA induces XRN2-mAID degradation. **d**, Schematic of the Pol III-inhibitor chase. Double-degron cells were pre-treated with AGB1 and 5-Ph-IAA for 48 h to deplete TRMT6 and XRN2 and to accumulate hypomodified pre- and mature tRNAᵢᴹᵉᵗ. At t = 0, 5-Ph-IAA was washed out (to allow XRN2 re-expression) and 30 µM ML-60218 was added (to inhibit new Pol III-driven transcription). Samples were collected at 0, 12, 24, 48, and 72 h post-washout. **e**, Western blot (upper panels) showing XRN2-mAID/XRN2, BromoTag-TRMT6/TRMT6, and β-tubulin levels, and Northern blot (lower panels) detecting pre-tRNAᵢᴹᵉᵗ and mature tRNAᵢᴹᵉᵗ across the chase. U6 snRNA, loading control. Lanes 1–2, baseline (no AGB1, no 5-Ph-IAA); Lanes 3, AGB1 +; Lane 4, AGB1 + 5-Ph-IAA pre-treatment at t = 0 (TRMT6 and XRN2 both depleted); lanes 5–8, AGB1 + ML-60218 + 5-Ph-IAA washout time course (TRMT6 depleted, XRN2 re-expressed); lanes 9–13, no AGB1 + ML-60218 + 5-Ph-IAA washout time course (BromoTag-TRMT6 intact, XRN2-mAID re-expressed). **e**, Quantification (lower panels) of mature tRNAᵢᴹᵉᵗ in e, normalized to U6 snRNA, displayed as relative expression versus chase time. Left, TRMT6-depleted condition (AGB1, lanes 4–8). Right, TRMT6-intact condition (no AGB1, lanes 9–13). Data are mean ± SD from three biological replicates.

To directly examine mature tRNA stability, we generated a dual-degron HCT116 line carrying BromoTag-TRMT6 together with an mAID (mini-AID) tag on the endogenous XRN2 C-terminus (Fig. 4c) ^32^. We pre-treated cells with AGB1 for 48 h to deplete TRMT6 and accumulate hypomodified mature tRNAᵢᴹᵉᵗ. Then, cells were shut down new Pol III-driven tRNA transcription with the Pol III inhibitor ML-60218 while concurrently inducing XRN2-mAID degradation with 5-Ph-IAA, and tracked the decay of the existing mature tRNAᵢᴹᵉᵗ pool over a 72 h chase (Fig. 4d). We pre-treated cells with AGB1 for 48 h to deplete TRMT6 and accumulate m^1^A-hypomodified precursor and mature tRNAᵢᴹᵉᵗ (Lane 4; Fig. 4e). We then shut down new Pol III-driven tRNA transcription with the Pol III inhibitor ML-60218 (30 µM) while concurrently inducing XRN2-mAID expression by washing out 5-Ph-IAA, and tracked the decay of the existing mature tRNAᵢᴹᵉᵗ pool over a 72 h chase (Lane 5-8; Fig. 4e). Northern blotting revealed that mature tRNAᵢᴹᵉᵗ in TRMT6-depleted cells progressively decreased during the chase following XRN2 recovery (Fig. 4e), whereas no apparent change of mature tRNAᵢᴹᵉᵗ was observed in TRMT6-expressing cells, indicating that XRN2 accounts for the majority of the observed decay of m¹A_58_-deficient mature tRNAᵢᴹᵉᵗ (Lane9-13; Fig. 4e). Together, these data indicate that m¹A_58_ loss generates two mechanistically distinct defects in human tRNAᵢᴹᵉᵗ metabolism: impaired precursor maturation and XRN2-mediated degradation of the mature hypomodified tRNA pool. Notably, the pre-tRNAᵢᴹᵉᵗ in TRMT6-depleted cells showed no change after XRN2 recovery (Lane4-9; Fig. 4e), indicating that hypomodified pre-tRNAᵢᴹᵉᵗ did not proceed the subsequent maturation process by RNase P and RNase Z.

### XRN2 co-depletion rescues the proliferation defect of TRMT6-depleted cells

To test if XRN2-mediated decay of mature tRNAᵢᴹᵉᵗ is functionally relevant for the cellular phenotype of TRMT6 loss, we monitored ATF4 expression and proliferation of BromoTag-TRMT6 cells (Fig. 5a, b). Consistent with the recovery of mature tRNAᵢᴹᵉᵗ level (Fig. 4a), XRN2 co-depletion decreased ATF4 expression (Fig. 5a) and restored proliferation in AGB1-treated BromoTag-TRMT6 cells (Fig. 5b), providing orthogonal support for the functional importance of XRN2-mediated control on cell proliferation.

**Fig. 5.**
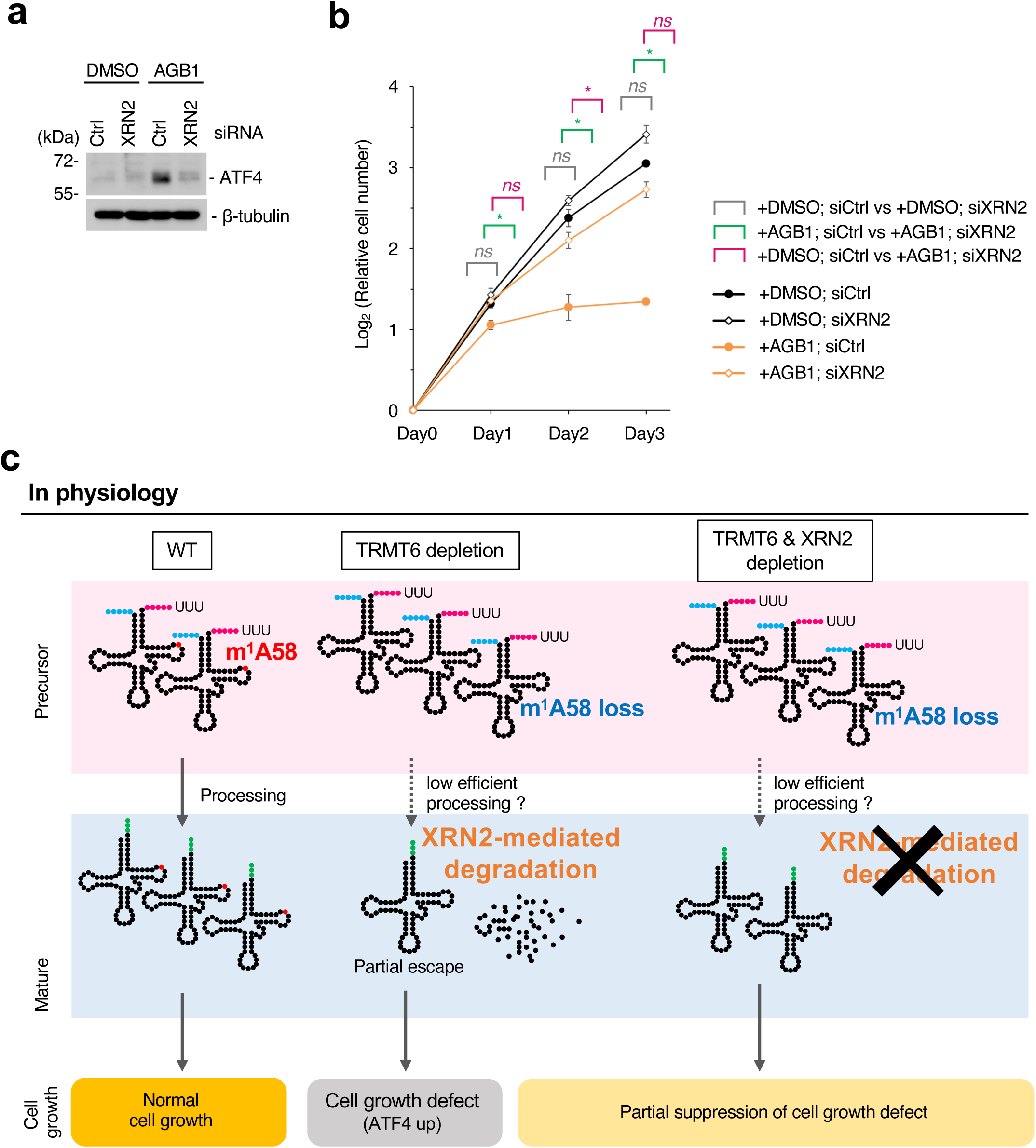
XRN2 co-depletion rescues the proliferation defect of TRMT6-depleted cells, and a two-step model for m¹A_58_-licensed tRNAᵢᴹᵉᵗ ribostasis. **a**, Western blot showing ATF4 and β-tubulin levels. **b**, Cell proliferation of BromoTag-TRMT6 KI#1 cells treated with DMSO or AGB1 (0.5 µM) in combination with siCtrl or siXRN2 (10 nM), over a 3-day time course. Cell number was counted at days 0–3. Data are mean ± SD from three biological replicates. Statistical significance was assessed using Student’s t-test. \**p* < 0.05. **c**, Model for the two-step regulation of tRNAᵢᴹᵉᵗ metabolism by m¹A_58_. Under physiological conditions (left), the TRMT6/TRMT61A complex installs m¹A_58_ on pre-tRNAᵢᴹᵉᵗ, supporting efficient maturation by RNase P and RNase Z and protecting mature tRNAᵢᴹᵉᵗ from XRN2-mediated decay. Upon TRMT6 depletion (middle), m¹A_58_ is lost; pre-tRNAᵢᴹᵉᵗ accumulates owing to low-efficiency processing, while the residual mature tRNAᵢᴹᵉᵗ pool is depleted by XRN2-mediated degradation. Upon co-depletion of TRMT6 and XRN2 (right), pre-tRNAᵢᴹᵉᵗ accumulation persists (XRN2-independent) but the mature tRNAᵢᴹᵉᵗ pool is partially preserved (XRN2-dependent).

## Discussion

Using an acute, ligand-inducible degron to deplete endogenous TRMT6 in human HCT116 cells, we show that m¹A_58_ maintains human tRNAᵢᴹᵉᵗ homeostasis through two separable pathways. First, m¹A_58_ promotes maturation of precursor tRNAᵢᴹᵉᵗ retaining 5′ leader and 3′ trailer sequences. Second, mature hypomodified tRNAᵢᴹᵉᵗ that escapes the maturation defect becomes susceptible to XRN2-mediated decay. Together, these pathways maintain the abundance and integrity of the mature tRNAᵢᴹᵉᵗ pool to sustain cell growth while avoiding excessive ISR activation (Fig. 5c).

The identification of a mammalian quality-control nuclease for hypomodified mature tRNAs has emerged only recently. We previously showed that XRN2, but not XRN1 or the nuclear-exosome catalytic subunit EXOSC10, constitutively degrades m⁷G-hypomodified mature tRNAs (tRNA^Val^-AAC and tRNA^Pro^) in HCT116 cells lacking METTL1 under physiological conditions, establishing the existence of an RTD-like activity in mammalian cells ^37^. This study extends the substrate scope of this activity to a structurally and functionally distinct class of hypomodified tRNAs — m¹A_58_-deficient mature tRNAᵢᴹᵉᵗ — and demonstrates that XRN2 acts on this second substrate. A key unresolved question is how XRN2 recognizes such structurally diverse hypomodified tRNAs. Because m⁷G_46_ and m¹A_58_ occupy distinct positions within the tRNA tertiary structure and are predicted to affect different structural features, our findings argue against a modification-specific recognition mechanism. Instead, they support a model in which loss of distinct tRNA modifications converges on a common structural consequence that renders mature tRNAs susceptible to XRN2-mediated decay. Indeed, previous studies in yeast proposed that hypomodified tRNAs become structurally destabilized and thereby expose normally protected RNA regions to exonucleolytic attack ^21,22^. In particular, elevated temperature is thought to exacerbate disruption of tRNA base-pairing interactions, promoting access of 5′→3′ exonucleases to single-stranded RNA regions ^23,24^. Our findings extend this concept in several important ways. First, XRN2-dependent decay occurs under physiological temperature conditions in mammalian cells. Second, the identification of both m⁷G-deficient elongator tRNAs and m¹A58-deficient initiator tRNAᵢᴹᵉᵗ as XRN2 substrates suggests that mammalian RTD-like surveillance monitors overall tRNA structural integrity rather than specific modification states.

m¹A_58_ has been proposed to stabilize the unique tertiary architecture of initiator tRNAᵢᴹᵉᵗ, including interactions that maintain the integrity of the tRNA core ^8,9^. Loss of this modification may therefore increase conformational flexibility or transient unfolding of mature tRNAᵢᴹᵉᵗ, potentially rendering the molecule more susceptible to XRN2-mediated decay. Alternatively, hypomodification may impair association of tRNAᵢᴹᵉᵗ with protective RNA-binding proteins or translation-initiation factors, indirectly exposing the tRNA to decay machinery. In our system, depletion of XRN2 largely restored tRNAᵢᴹᵉᵗ abundance as well as downstream phenotypes, supporting the idea that XRN2-mediated decay is a major consequence of TRMT6 loss (Fig. 5a, b). Nevertheless, the known structural role of m¹A_58_ suggests that this modification may contribute more broadly to the stability and functional integrity of tRNAᵢᴹᵉᵗ. Although our data support an important contribution of XRN2-mediated tRNAᵢᴹᵉᵗ decay, we cannot exclude the possibility that additional tRNA species or other downstream effects of TRMT6 loss also contribute to the observed phenotypes.

In addition to its role in protecting mature tRNAᵢᴹᵉᵗ from decay, our findings reveal an unexpected requirement for m¹A_58_ in precursor tRNA maturation. TRMT6 depletion resulted in the accumulation of immature tRNAᵢᴹᵉᵗ species retaining 5′ leader and 3′ trailer sequences (Fig. 2e; Supplementary Fig. 3a, b), indicating that m¹A_58_ promotes efficient processing during early stages of tRNA biogenesis. In budding yeast, defective pre-tRNAᵢᴹᵉᵗ can be targeted by the TRAMP complex and nuclear exosome through oligo(A)-mediated surveillance pathways ^19,20,38,39^. However, despite the accumulation of immature tRNAᵢᴹᵉᵗ species upon TRMT6 depletion, we did not detect clear evidence of polyadenylated pre-tRNAᵢᴹᵉᵗ intermediates under our experimental conditions (data not shown). These observations suggest that m¹A_58_-deficient pre-tRNAᵢᴹᵉᵗ is not efficiently routed into canonical TRAMP-dependent nuclear RNA surveillance pathways, at least as a major fate. A recent independent study by Liu et al. reaches a convergent conclusion that TRMT6/TRMT61A-mediated m¹A_58_ supports human pre-tRNA maturation and protects tRNAᵢᴹᵉᵗ from XRN2-dependent surveillance. Notably, Liu et al. also reported that depletion of the nuclear exosome component EXOSC10/RRP6 did not lead to pre-tRNAᵢᴹᵉᵗ accumulation in mammalian cells, consistent with our interpretation that m¹A_58_-deficient pre-tRNAᵢᴹᵉᵗ primarily reflects impaired processing rather than nuclear exosome-mediated decay^40^. Together, these observations support a model in which the primary consequence of m¹A_58_ deficiency is impaired pre-tRNA processing rather than active nuclear exosome-mediated degradation.

Our findings also raise the question of where XRN2-mediated surveillance occurs within the cell. Because XRN2 is predominantly nuclear, hypomodified mature tRNAᵢᴹᵉᵗ may undergo nuclear retention or re-import prior to degradation. Such a mechanism would parallel nuclear surveillance pathways proposed in yeast, in which defective mature tRNAs undergo retrograde transport to the nucleus and are selectively targeted for repair or decay ^41–43^. Determining how hypomodified mature tRNAs encounter XRN2 will therefore be important for understanding the spatial organization of mammalian tRNA quality control.

Collectively, our study identifies m¹A_58_ as a critical determinant of human initiator tRNA homeostasis (ribostasis) and establishes that mammalian cells employ multiple surveillance mechanisms to eliminate defective tRNAᵢᴹᵉᵗ species at distinct stages of the tRNA life cycle. These findings further support the existence of a conserved RTD-like pathway in mammalian cells and suggest that mammalian cells employ layered quality-control mechanisms to maintain initiator tRNA integrity throughout the tRNA life cycle.

## Methods

### Cell lines

HCT116 cells (human colorectal carcinoma; ATCC, CCL-247) were cultured as described previously ^37^. All HCT116 cell lines expressing degron-fused proteins were generated as previously described ^32,37^. To generate BromoTag-TRMT6 knock-in cells, HCT116 cells were co-transfected with the BSD-P2A-BromoTag donor plasmid and the pX330-sgRNA-TRMT6 plasmid targeting the N-terminus of TRMT6, selected with blasticidin S at 6.25 ng/µL, and screened by genomic PCR. Two independent BromoTag-TRMT6 N-terminal knock-in clones (KI#1 and KI#2) were used in parallel throughout this study. The XRN2-mAID double-degron line was generated by sequential CRISPR knock-in of the pBS-mAID-Hygro donor at the C-terminus of XRN2 on one of the BromoTag-TRMT6 N-terminal knock-in clones in the HCT116 OsTIR1(F74G) background, using procedures described previously ^32^.

### Plasmid constructions

The CRISPR-Cas9 backbone plasmid ^44^, the BSD-P2A-BromoTag donor ^32^, and the pBS-mAID-Hygro donor ^45^ were used as the backbone constructs. Plasmids generated in this study include the pX330 vector encoding the sgRNA targeting the N-terminus of human TRMT6 (sgRNA-TRMT6: 5′-CGACCGGCTGAGCGTCATGG/AGG-3′) and the corresponding BSD-P2A-BromoTag donor with ∼500-bp homology arms flanking the TRMT6 start codon. The pX330 vector encoding the sgRNA targeting the C-terminus of human XRN2 (sgRNA-XRN2: 5′-TGTAGAATGATGAAAGGATT/TGG-3′) and the pBS-XRN2-mAID-Hygro donor were used. All constructs generated in this study were verified by Sanger sequencing.

### Antibodies

For immunoblotting, the following primary antibodies were used: anti-TRMT6 (rabbit polyclonal, ProteinTech, 16727-1-AP), anti-ATF4 (rabbit monoclonal, Cell Signaling Technology, #11815, clone D4B8), anti-XRN2 (rabbit polyclonal, ProteinTech, 112671-1-AP), and anti-β-tubulin (mouse monoclonal, Developmental Studies Hybridoma Bank, E7). HRP-conjugated anti-rabbit IgG (Cell Signaling Technology, 7074) and HRP-conjugated anti-mouse IgG (Thermo Fisher Scientific, 32230) were used as secondary antibodies. For RNA immunodetection of N¹-methyladenosine (m¹A), an anti-m¹A monoclonal antibody (clone D345-3; MBL International; catalog number to be supplied) was used as the primary antibody.

### Degrader compounds and Pol III inhibitor

AGB1 (Tocris, 7686) was used at 0.5 µM; 5-Ph-IAA was synthesized as previously described ^46^ and used at 1 µM; the Pol III inhibitor ML-60218 (MedChemExpress, HY-122122) ^47^ was used at 30 µM. Detailed properties and handling of these compounds are as described previously ^37^.

### Inducible degradation of TRMT6

For inducible degradation of BromoTag-TRMT6, BromoTag-TRMT6 cells (KI#1 or KI#2) were treated with 0.5 µM AGB1 for the indicated times. Near-complete depletion of BromoTag-TRMT6 protein was achieved within 12 h of treatment (Fig. 1b).

### Combined BromoTag-mediated TRMT6 degradation and siRNA-mediated XRN2 knockdown

For combined BromoTag-mediated degradation of TRMT6 and siRNA-mediated knockdown of XRN2, BromoTag-TRMT6 cells (KI#1) were seeded at 1.5 × 10⁵ cells per well in 6-well plates. After 24 h, cells were treated with 0.5 µM AGB1 and simultaneously transfected with siRNA against XRN2 (siXRN2) or non-targeting control siRNA (siCtrl; MISSION siRNA Universal Negative Control #1, Sigma-Aldrich, SIC001) at a final concentration of 10 nM using Lipofectamine RNAiMAX (Thermo Fisher Scientific, 13778-075) as described previously ^37^. Cells were harvested 72 h after the start of treatment for RNA and protein analyses. The siRNA sequences are listed in Supplementary Table 1.

### XRN2-mAID double-degron and Pol III-inhibitor chase

The Pol III-inhibitor chase using the XRN2-mAID degron and ML-60218 was performed as described previously ^37^ with the following modifications for the TRMT6 context. Double-degron (BromoTag-TRMT6; XRN2-mAID) cells were pre-treated with 0.5 µM AGB1 and 1 µM 5-Ph-IAA for 48 h to deplete TRMT6 and XRN2 and to accumulate hypomodified pre- and mature tRNAᵢᴹᵉᵗ. At the start of the chase (t = 0), 5-Ph-IAA was removed by two PBS washes followed by replacement with fresh ligand-free medium to allow XRN2 re-expression, and 30 µM ML-60218 was added to inhibit new Pol III-driven tRNA transcription; AGB1 was maintained throughout the chase to keep TRMT6 depleted. Cells were sampled at 0, 12, 24, 48, and 72 h post-washout for Northern and Western blot analysis. Control conditions in which TRMT6 was not depleted (no AGB1) were processed in parallel.

### siRNA transfection

siRNA transfections were performed as described previously ^37^. Briefly, HCT116-derived cells were seeded at 1.0–1.5 × 10⁵ cells per well in 6-well plates 24 h before transfection, transfected with siRNAs at 10 nM using Lipofectamine RNAiMAX (Thermo Fisher Scientific, 13778-075) in Opti-MEM (Thermo Fisher Scientific, 31985-070), and harvested 48 or 72 h after transfection as specified in the figure legends. MISSION siRNA Universal Negative Control #1 (Sigma-Aldrich, SIC001) was used as the non-targeting control. siRNA sequences are listed in Supplementary Table 1.

### Plasmid transfection

Plasmid transfections were performed as described previously ^37^. Briefly, HCT116-derived cells were seeded at 1.0 × 10⁵ cells per well in 6-well plates 24 h before transfection, transfected with 25 ng of plasmid DNA using ViaFect Transfection Reagent (Promega, E4981), and harvested at 72 h.

### RNA extraction

Total RNA was extracted with ISOGEN (Nippon Gene, 319-90211) as described previously ^37,48^.

### Northern blotting

Northern blotting was performed as described previously ^37,48^ with the addition of fluorescent-probe detection. In brief, 1–2 µg of total RNA was resolved on 12% urea–PAGE gels (7 M urea, 1× TBE, 12% acrylamide), transferred onto Hybond N+ membranes (Cytiva), UV-crosslinked, and hybridized at 42 °C overnight with 3′-DIG-labeled DNA probes (Merck, 3353575910) or with DY800- or Cy5-labeled fluorescent DNA probes (50 pmol per membrane). All DNA probes were synthesized by Eurofins Genomics (Tokyo, Japan); DIG-labeled probes were used to detect tRNA^Lys^-TTT and tRNA^Tyr^-GTA, a Cy5-labeled probe was used to detect tRNAᵢᴹᵉᵗ, U6, and a DY800-labeled probe was used to detect U6 snRNA as the loading control. DIG-probe signals were developed with CDP-Star (Merck, 12041677001) following anti-DIG-AP incubation (Merck, 11093274910); fluorescent signals from DY800 and Cy5 were detected directly using a ChemiDoc Touch imaging system (Bio-Rad). Band intensities were quantified using Image Lab software (Bio-Rad) and tRNA signals were normalized to the U6 snRNA signal on the same membrane unless otherwise noted. DNA probe sequences are listed in Supplementary Table 2.

### Anti-m¹A RNA immunodetection

After standard urea–PAGE transfer to Hybond N+ membrane and UV-crosslinking, the membrane was blocked with 5% non-fat dry milk in PBS and incubated 25 °C with anti-m¹A monoclonal antibody (clone D345-3; MBL International) diluted in PBS-T. After PBS-T washes, the membrane was incubated with HRP-conjugated anti-mouse IgG (Thermo Fisher Scientific, 32230) and developed with ECL substrate (Amersham) on a ChemiDoc Touch imaging system. The membrane was subsequently stripped with Restore Western Blot Stripping Buffer (Thermo Fisher Scientific) and re-hybridized with a DY800-labeled U6 snRNA probe as a loading control as described under Northern blotting.

### Western blotting

Western blotting was performed as described previously ^37,48^. Briefly, cells were lysed in 2 × SDS sample buffer, resolved by 6–12% SDS-PAGE, transferred to nitrocellulose membrane (Amersham), blocked with 5% skim milk in PBS(-) (Nacalai Tesque), and incubated with primary antibodies (see Antibodies) and HRP-conjugated secondary antibodies. Signals were visualized with ECL Prime or ECL Select Western Blotting Detection Reagent (Amersham) and imaged with a ChemiDoc Touch imaging system (Bio-Rad). Band intensities were quantified using Image Lab software (Bio-Rad).

### RNA-seq sample preparation and library construction

Sample preparation and RNA-seq library construction were performed as described previously ^37,48^. Briefly, total RNA was treated with DNase I (NEB) and cleaned up with the NucleoSpin RNA Clean-up XS Kit (MACHEREY-NAGEL); the small-RNA fraction (<200 nt) was then enriched with the mirVana miRNA Isolation Kit (Thermo Fisher Scientific, AM1561) and treated with recombinant E. coli AlkB and AlkB-D135S to remove RT-blocking methylations. Demethylated RNAs were used as input for library construction with the NEBNext Multiplex Small RNA Library Prep Set for Illumina (NEB, E7300S) using Maxima H Minus Reverse Transcriptase (Thermo Fisher Scientific, EP0752) in place of the kit-provided reverse transcriptase. Libraries were sequenced on a NovaSeq X Plus instrument (Illumina) in paired-end 150-bp mode. Biological triplicates were sequenced for each cell line (BromoTag-TRMT6 KI#1 and KI#2) and each condition (DMSO or AGB1).

### Quantification and statistical analysis

RNA-seq read processing and differential expression analysis were performed as described previously ^37,48^. Briefly, raw paired-end reads were processed through the tRAX pipeline to merge them into single-end reads, trim adapters, and map to a reference sequences built from 432 mature human tRNA sequences ^49^ together with other non-coding small RNAs. Reads derived from mature tRNAs and from precursor tRNAs (containing 5′-leader and/or 3′-trailer sequences) were separately quantified using BWA v0.7.17 ^50^ followed by TIGAR2 ^51^ and differentially expressed genes were extracted using edgeR v3.36.0 ^52^ at adjusted *P* < 0.05 (Benjamini–Hochberg) and |log₂ fold-change| ≥ 1. Northern and Western blot band intensities were quantified with Image Lab software (Bio-Rad) and normalized to U6 snRNA on the same membrane. Condition means were compared by an unpaired, two-tailed Welch′s t-test; two-way ANOVA (TRMT6 status × XRN2 status) was applied where appropriate, and pairwise post-hoc comparisons were Bonferroni-corrected as noted in the figure legends. *P* < 0.05 was considered significant. All quantitative experiments were performed with at least three independent biological replicates unless otherwise stated.

### Cell proliferation assay

Cells were seeded at 1 × 10^5^ cells per well in 6-well plates 24 h before the start of the time course. At day 0, cells were treated with 0.5 µM AGB1, 10 nM siRNA, or the indicated combination. Cell numbers were counted at days 0–3 using an automated cell counter (Bio-Rad TC20) in triplicate biological replicates.

## Data and code availability

The RNA-seq data of small RNAs have been deposited with links to BioProject accession number PRJDB42361 in the DDBJ BioProject database.

## Acknowledgments

We are grateful to T. Miyoshi for technical assistance. We also thank Dr. Takeshi Chujo (Kumamoto University) for technical advice and helpful discussions during manuscript preparation. This work was supported by JSPS KAKENHI grants JP25K09570 (K.M.), JP22H02669 (K.S.), JP23K23932 (K.S.), and JP25K02209 (K.S.), as well as by the Takeda Science Foundation (K.S.).

## Author contributions

Kiito Otsubo: investigation; formal analysis; methodology; validation; writing-original draft; writing-review and editing. Mio Nagura: investigation; methodology; validation; writing-original draft; writing-review and editing. Makoto Terauchi: data curation; software; investigation; methodology and writing-original draft. Hideki Noguchi: data curation; software; methodology; and writing-review. Masato T. Kanemaki: methodology and editing. Keita Miyoshi: investigation; methodology; validation; writing-review and editing. Kuniaki Saito: conceptualization; funding acquisition; investigation; methodology; project administration; supervision; writing-original draft; writing-review and editing.

## Supplementary figure legends

**Supplementary Fig. 1.**
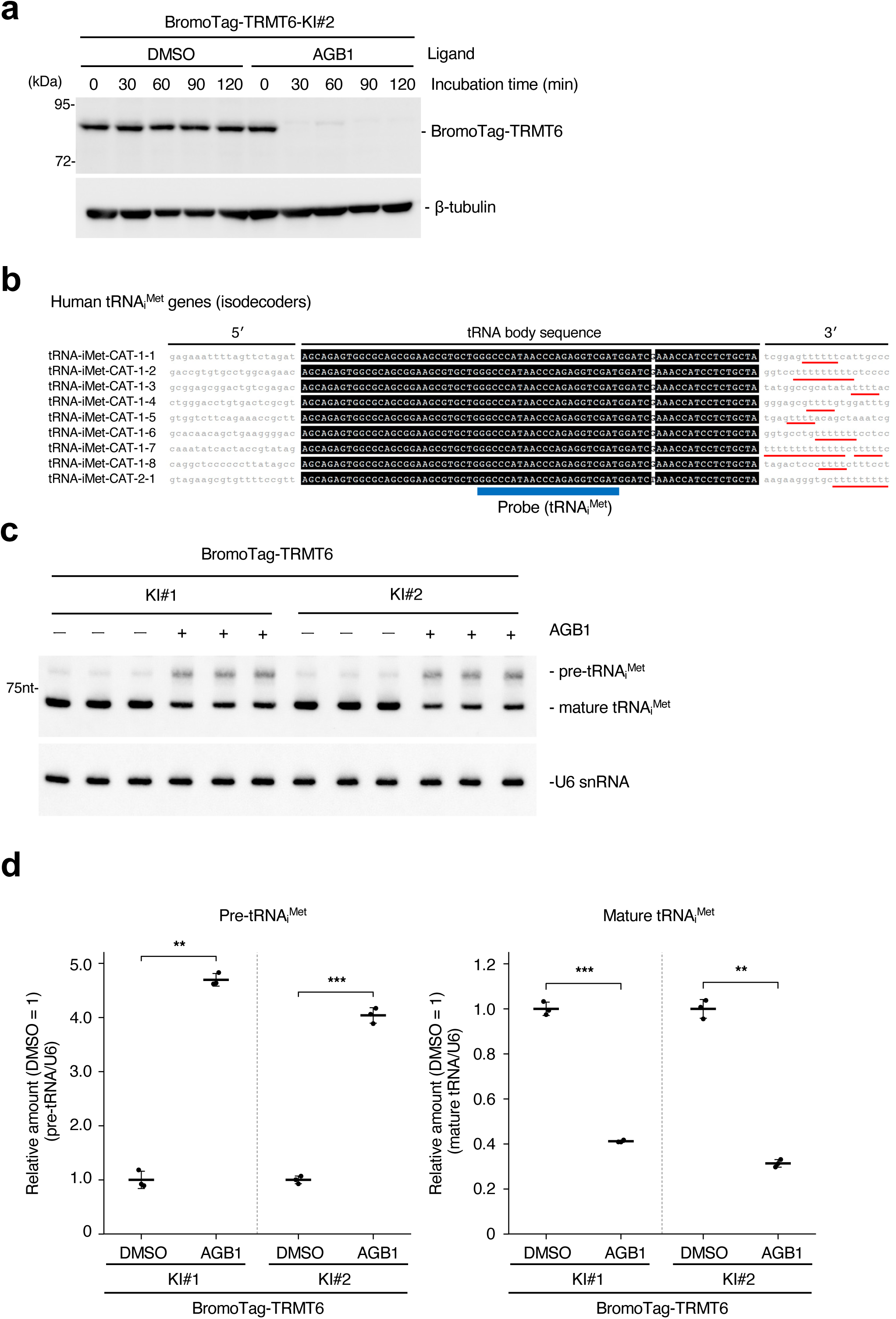
Probe design used for tRNAᵢᴹᵉᵗ Northern blotting for precursor and mature tRNAᵢᴹᵉᵗ in TRMT6-depleted cells. **a**, Western blot showing AGB1 (0.5 µM)-induced degradation of BromoTag-TRMT6 in HCT116 (KI#2) over a 0–120 min time course. β-tubulin, loading control. **b**, Schematic of the DIG-labeled oligonucleotide probe complementary to the conserved tRNA body region shared among all tRNAᵢᴹᵉᵗ isodecoders. Probe sequence is listed in Supplementary Table 2. **c**, Northern blot of total RNA from BromoTag-TRMT6 cells treated with DMSO or AGB1 for 48 h as in Fig. 1e, probed with oligonucleotide complementary to the conserved body region of tRNAᵢᴹᵉᵗ (Supplementary Fig. 1b). **d**, Relative amounts of pre-iMet and mature-iMet in Supplementary Fig. 1b. Data are presented as mean ± SD with individual biological replicates shown as dots (n = 3). Statistical significance was assessed using two-tailed Student’s t-test. **p < 0.01, ***p < 0.001 vs. DMSO.

**Supplementary Fig. 2.**
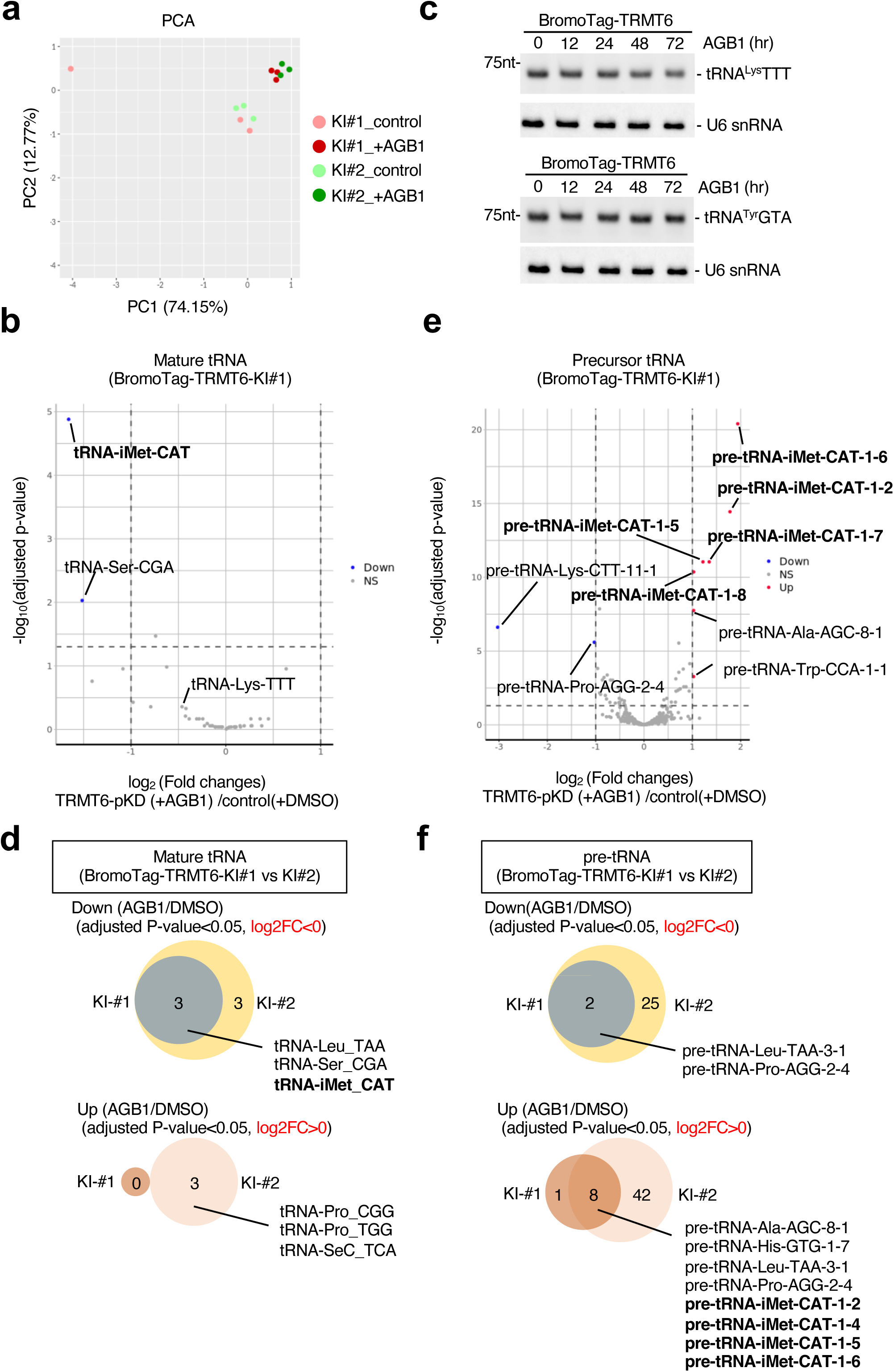
RNA-seq and Northern blot analyses in TRMT6-depleted cells. **a**, Principal-component analysis (PCA) of tRNA-seq libraries from KI#1 and KI#2 cells treated with DMSO or AGB1, showing clean separation of AGB1 from DMSO on PC1 across both clones and tight clustering of biological replicates. **b**, Volcano plot of tRNA-seq differential expression at the mature-pool, anticodon level in KI#1 cells treated with AGB1 versus DMSO. tRNAs with adjusted *P*-value < 0.05 and |log₂ fold change| ≥ 1 were considered differentially expressed. **c**, Northern blot of representative non-iMet tRNAs (tRNA^Lys^-TTT and tRNA^Tyr^-GTA) in BromoTag-TRMT6 cells treated with AGB1 over the indicated time course, showing no substantial change with TRMT6 depletion. U6 snRNA, loading control. **d**, Intersection of significantly down-regulated mature tRNAs between KI#1 and KI#2 (adjusted *P* < 0.05, log₂FC ≤ −1) as in main Fig. 2C, displayed for the KI#1 dataset. **e**, Volcano plot of tRNA-seq differential expression for the precursor read population in KI#1 cells. Significance and labeling thresholds as in b.

**Supplementary Fig. 3.**
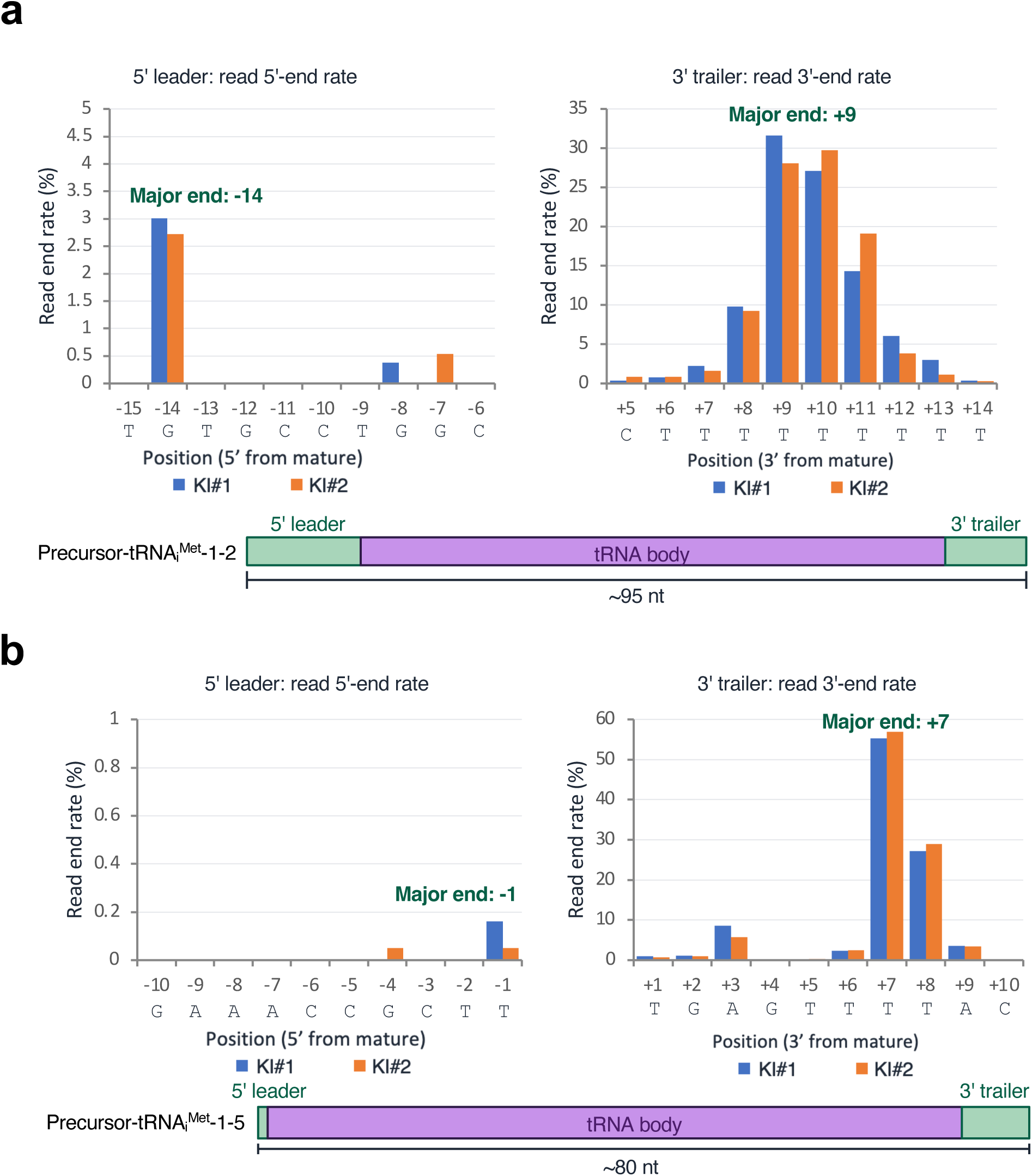
Read-end mapping for additional tRNAᵢᴹᵉᵗ-CAT isodecoders. **a**, Read 5′-end and 3′-end mapping for pre-tRNAᵢᴹᵉᵗ-CAT-1-2 in TRMT6-depleted cells, calculated as in main Fig. 2E from three biological replicates. Green boxes mark positions with the highest frequencies. The schematic at the bottom illustrates the inferred pre-tRNAᵢᴹᵉᵗ-CAT-1-2 structure with retained 5′-leader and 3′-trailer sequences. **b**, As in a, for pre-tRNAᵢᴹᵉᵗ-CAT-1-5. The major end positions differ between isodecoders, reflecting differences in their genomic context.

**Supplementary Table 1.**
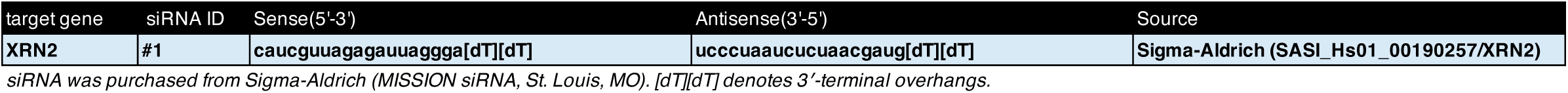
List of siRNAs used in this study.

**Supplementary Table 2.**
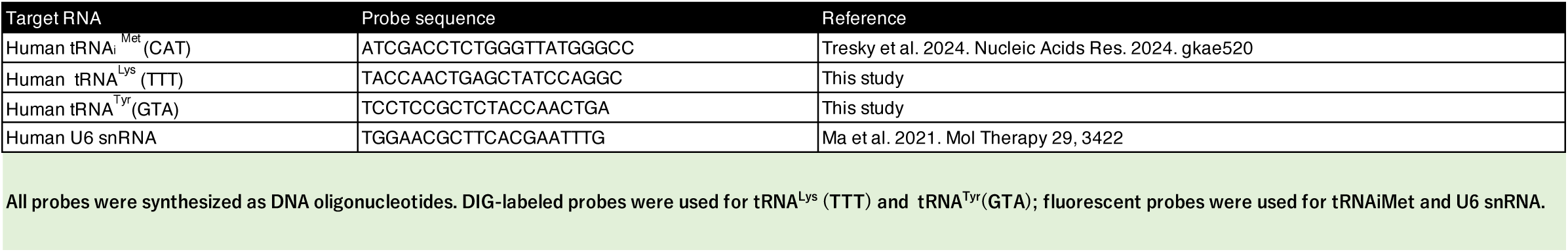
List of DNA probes used in this study.

## References

1. Boccaletto, P. & et al. MODOMICS: a database of RNA modification pathways. 2021 update. Nucleic Acids Res 50, D231–D235 (2022).

2. Motorin, Y. & Helm, M. RNA nucleotide methylation. Wiley Interdiscip. Rev. RNA 2, 611–631 (2011).

3. Pan, T. Modifications and functional genomics of human transfer RNA. Cell Res 28, 395–404 (2018).

4. Roundtree, I.A., Evans, M.E., Pan, T. & He, C. Dynamic RNA modifications in gene expression regulation. Cell 169, 1187–1200 (2017).

5. Suzuki, T. The expanding world of tRNA modifications and their disease relevance. Nat. Rev. Mol. Cell Biol 22, 375–392 (2021).

6. Orellana, E.A., Siegal, E. & Gregory, R.I. tRNA dysregulation and disease. Nat. Rev. Genet 23, 651–664 (2022).

7. Torres, A.G., Batlle, E. & Pouplana, L. Role of tRNA modifications in human diseases. Trends Mol. Med 20, 306–314 (2014).

8. Anderson, J. & et al. The essential Gcd10p–Gcd14p nuclear complex is required for 1-methyladenosine modification and maturation of initiator methionyl-tRNA. Genes Dev 12, 3650–3662 (1998).

9. Ozanick, S., Krecic, A., Andersland, J. & Anderson, J.T. The bipartite structure of the tRNA m¹A58 methyltransferase from S. cerevisiae is conserved in humans. RNA 11, 1281–1290 (2005).

10. Anderson, J.T. & Droogmans, L. Biosynthesis and function of 1-methyladenosine in transfer RNA. Vol. 12 121–139 (2005).

11. Finer-Moore, J., Czudnochowski, N., Connell, J.D., Wang, A.L. & Stroud, R.M. Crystal structure of the human tRNA m¹A58 methyltransferase–tRNA³^Lys complex: refolding of substrate tRNA allows access to the methylation target. J. Mol. Biol 427, 3862–3876 (2015).

12. Helm, M., Giegé, R. & Florentz, C. A Watson-Crick base-pair-disrupting methyl group (m1A9) is sufficient for cloverleaf folding of human mitochondrial tRNALys. Biochemistry 38, 13338–13346 (1999).

13. Oliva, R., Cavallo, L. & Tramontano, A. Accurate energies of hydrogen bonded nucleic acid base pairs and triplets in tRNA tertiary interactions. Nucleic Acids Res 34, 865–879 (2006).

14. Kleiber, N. et al. The RNA methyltransferase TRMT6/TRMT61A targets diverse RNA species and beyond. Nucleic Acids Res 50, 9444–9461 (2022).

15. Costa-Mattioli, M. & Walter, P. The integrated stress response: from mechanism to disease. Science 368, eaat5314 (2020).

16. Pavitt, G.D. Regulation of translation initiation factor eIF2B at the hub of the integrated stress response. Wiley Interdiscip. Rev. RNA 9, e1491 (2018).

17. Sonenberg, N. & Hinnebusch, A.G. Regulation of translation initiation in eukaryotes: mechanisms and biological targets. Cell 136, 731–745 (2009).

18. Calvo, O. et al. GCD14p, a repressor of GCN4 translation, cooperates with Gcd10p and Lhp1p in the maturation of initiator methionyl-tRNA in Saccharomyces cerevisiae. Mol. Cell. Biol 19, 4167–4181 (1999).

19. Kadaba, S. & et al. Nuclear surveillance and degradation of hypomodified initiator tRNAMet in S. cerevisiae. Genes Dev 18, 1227–1240 (2004).

20. LaCava, J. & et al. RNA degradation by the exosome is promoted by a nuclear polyadenylation complex. Cell 121, 713–724 (2005).

21. Alexandrov, A. & et al. Rapid tRNA decay can result from lack of nonessential modifications. Mol. Cell 21, 87–96 (2006).

22. Chernyakov, I., Whipple, J.M., Kotelawala, L., Grayhack, E.J. & Phizicky, E.M. Degradation of several hypomodified mature tRNA species in Saccharomyces cerevisiae is mediated by Met22 and the 5&#x2019;-3&#x2019; exonucleases Rat1 and Xrn1. Genes Dev 22, 1369–1380 (2008).

23. Dewe, J.M., Whipple, J.M., Chernyakov, I., Jaramillo, L.N. & Phizicky, E.M. The yeast rapid tRNA decay pathway competes with elongation factor 1A for substrate tRNAs and acts on tRNAs lacking one or more of several modifications. RNA 18, 1886–1896 (2012).

24. Whipple, J.M., Lane, E.A., Chernyakov, I., Silva, S. & Phizicky, E.M. The yeast rapid tRNA decay pathway primarily monitors the structural integrity of the acceptor and T-stems of mature tRNA. Genes Dev 25, 1173–1184 (2011).

25. Tasak, M. & Phizicky, E.M. Initiator tRNA lacking 1-methyladenosine is targeted by the rapid tRNA decay pathway in evolutionarily distant yeast species. PLoS Genet 18, e1010215 (2022).

26. Ding, Y. et al. TRMT6/TRMT61A-mediated tRNA m(1)A modification enhances protein translation and activates the IRE1alpha-XBP1s pathway to promote anaplastic thyroid cancer progression. Cell Mol Biol Lett 31(2026).

27. Monshaugen, I. et al. Depletion of the m1A writer TRMT6/TRMT61A reduces proliferation and resistance against cellular stress in bladder cancer. Front Oncol 13, 1334112 (2023).

28. Tao, E.W. et al. TRMT6-mediated tRNA m1A modification acts as a translational checkpoint of histone synthesis and facilitates colorectal cancer progression. *Nat*. Cancer 6, 1458–1476 (2025).

29. Wang, Y. & et al. N¹-methyladenosine methylation in tRNA drives liver tumourigenesis by regulating cholesterol metabolism. Nat. Commun 12, 6314 (2021).

30. Macari, F. & et al. TRM6/61A-dependent base methylation of tRNA-derived fragments regulates gene-silencing activity and the unfolded protein response in bladder cancer. Oncotarget 7, 50943–50954 (2016).

31. Watanabe, K. et al. Degradation of initiator tRNAMet by Xrn1/2 via its accumulation in the nucleus of heat-treated HeLa cells. Nucleic Acids Res 41, 4671–4685 (2013).

32. Hatoyama, Y. & et al. Combination of AID2 and BromoTag expands the utility of degron-based protein knockdowns. EMBO Rep 25, 4062–4077 (2024).

33. Bond, A.G. & et al. Development of BromoTag: A &#x201C;bump-and-hole&#x201D;-PROTAC system to induce potent, rapid, and selective degradation of tagged target proteins. J. Med. Chem 64, 15477–15502 (2021).

34. Fukuda, H. et al. Cooperative methylation of human tRNA3Lys at positions A58 and U54 drives the early and late steps of HIV-1 replication. Nucleic Acids Res 49, 11855–11867 (2021).

35. Safra, M. et al. The m1A landscape on cytosolic and mitochondrial mRNA at single-base resolution. Nature 551, 251–255 (2017).

36. Hinnebusch, A.G. The scanning mechanism of eukaryotic translation initiation. Annu. Rev. Biochem 83, 779–812 (2014).

37. Miyoshi, K. et al. The mammalian rapid tRNA decay pathway is critical for N⁷-methylguanosine-hypomodified tRNA degradation under physiological conditions. bioRxiv (2025).

38. Vanácová, S. & et al. A new yeast poly(A) polymerase complex involved in RNA quality control. PLoS Biol 3, e189 (2005).

39. Wang, X., Jia, H., Jankowsky, E. & Anderson, J.T. Degradation of hypomodified tRNA(iMet) in vivo involves RNA-dependent ATPase activity of the DExH helicase Mtr4p. RNA 14, 107–116 (2008).

40. Liu, X. et al. TRMT6/61A-mediated m¹A methylation facilitates human pre-tRNA maturation and prevents surveillance by XRN2. bioRxiv (2026).

41. Shaheen, H.H. & Hopper, A.K. Retrograde movement of tRNAs from the cytoplasm to the nucleus in Saccharomyces cerevisiae. Proc. Natl. Acad. Sci. USA 102, 11290–11295 (2005).

42. Takano, A., Endo, T. & Yoshihisa, T. tRNA actively shuttles between the nucleus and cytosol in yeast. Science 309, 140–2 (2005).

43. Whitney, M.L., Hurto, R.L., Shaheen, H.H. & Hopper, A.K. Rapid and reversible nuclear accumulation of cytoplasmic tRNA in response to nutrient availability. Mol. Biol. Cell 18, 2678–2686 (2007).

44. Cong, L. & et al. Multiplex genome engineering using CRISPR/Cas systems. Science 339, 819–823 (2013).

45. Natsume, T., Kiyomitsu, T., Saga, Y. & Kanemaki, M.T. Rapid protein depletion in human cells by auxin-inducible degron tagging with short homology donors. Cell Rep 15, 210–218 (2016).

46. Yesbolatova, A. & et al. The auxin-inducible degron 2 technology provides sharp degradation control in yeast, mammalian cells, and mice. Nat. Commun 11, 5701 (2020).

47. Wu, L. et al. Novel small-molecule inhibitors of RNA polymerase III. Eukaryot. Cell 2, 256–264 (2003).

48. Kaneko, S. & et al. Mettl1-dependent m7G tRNA modification is essential for maintaining spermatogenesis and fertility in Drosophila melanogaster. Nat. Commun 15, 8147 (2024).

49. Chan, P.P. & Lowe, T.M. GtRNAdb: a database of transfer RNA genes detected in genomic sequence. Nucleic Acids Res 37, D93–7 (2009).

50. Li, H. & Durbin, R. Fast and accurate short read alignment with Burrows-Wheeler transform. Bioinformatics 25, 1754–1760 (2009).

51. Nariai, N. & et al. TIGAR2: sensitive and accurate estimation of transcript isoform expression with longer RNA-Seq reads. BMC Genomics 15 Suppl 10, S5 (2014).

52. Robinson, M.D., McCarthy, D.J. & Smyth, G.K. edgeR: a Bioconductor package for differential expression analysis of digital gene expression data. Bioinformatics 26, 139–140 (2010).

